# IL-26 from ILCs regulates early-life gut epithelial homeostasis by shaping microbiota composition

**DOI:** 10.1101/2025.05.05.652124

**Authors:** Yazan Salloum, Gwendoline Gros, Keinis Quintero-Castillo, Camila Garcia-Baudino, Soraya Rabahi, Akshai Janardhana Kurup, Patricia Diabangouaya, David Pérez-Pascual, Rodrigo A. Morales Castro, Jos Boekhorst, Eduardo J. Villablanca, Jean-Marc Ghigo, Carmen G. Feijoo, Sylvia Brugman, Pedro P. Hernandez

**Affiliations:** Institut Curie, PSL Research University CNRS UMR 3215, INSERM U934, 26 Rue d’Ulm, 75248 Paris Cedex 05, France; Fish Immunology Laboratory, Faculty of Life Science, Andres Bello University, Santiago 8370146, Chile; Institut Pasteur, Université Paris-Cité, UMR CNRS 6047, Genetics of Biofilms Laboratory, Department of Microbiology, Paris, France; Department of Medicine Solna (MedS), Karolinska Institutet, and Division of Immunology and Respiratory Medicine, Karolinska University Hospital, Solna, and Center for Molecular Medicine, Karolinska University Hospital, SE-171 76, Stockholm, Sweden; Division of Clinical Immunology, Department of Laboratory Medicine (Labmed). Karolinska Institutet. 141 52 Huddinge, Sweden; Host-Microbe Interactomics, Animal Sciences Group, Wageningen University & Research, Wageningen, Netherlands

**Author notes:** Corresponding author. (PPH).

## Abstract

Animals host symbiotic microbial communities that shape gut health. However, how the host immune system and microbiota interact to regulate epithelial homeostasis, particularly during early development, remains unclear. Human interleukin-26 (IL-26) is associated with gut inflammation and has intrinsic bactericidal activity *in vitro*, yet its *in vivo* functions are largely unknown, primarily due to its absence in rodents. To examine the role of IL-26 in early life, we used zebrafish and found that gut epithelial cells in *il26^-/-^* larvae exhibited increased proliferation and DNA damage, faster turnover, and an altered abundance of cell populations. This epithelial dysregulation occurred independently of the IL-26 canonical receptor and resulted from dysbiosis in *il26^-/-^*. Moreover, IL-26 bactericidal activity was conserved in zebrafish. Our findings suggest that IL-26 directly regulates microbiota composition through this property. We further identified innate lymphoid cells (ILCs) as the primary source of IL-26 at this developmental stage. These findings establish IL-26 as a central player in a regulatory circuit linking the microbiota, ILCs, and intestinal epithelial cells to maintain gut homeostasis during early life.

## INTRODUCTION

The gastrointestinal tract is a complex system housing numerous populations of immune cells and commensal microorganisms. However, it is also a port of entry for various pathogens. During early life, a dynamic crosstalk between immune cells, epithelial cells, and colonizing microbiota is critical for establishing life-long intestinal homeostasis (*1–3*). Dysregulation of this crosstalk can lead to inflammatory bowel disease (IBD), a group of disorders closely associated with alterations in the gut microbiota that can elicit a chronic immune response and lead to tissue damage. In addition, IBD patients are at higher risk of developing colon cancer. A hallmark of this cancer is greater genomic instability and increased proliferation of abnormal cells (*4*). Therefore, identifying factors that regulate early-life microbiota composition, epithelial cell proliferation, and DNA damage is pivotal for advancing early diagnosis, prevention, and treatment of these pathologies.

Genome-wide association studies have identified interleukin-26 (IL-26) as a risk locus for ulcerative colitis (UC) (*5*), indicating a potential role of this cytokine in intestinal homeostasis. Moreover, IL-26 has been found to be overexpressed in the inflamed mucosa of IBD patients (*6*, *7*). Notably, intestinal epithelial cells (IECs) were shown to express the IL-26 receptor complex (IL10RB and IL20RA) and to respond to this cytokine *in vitro* (*7*). Despite these findings, the *in vivo* functions of IL-26 and its impact on IECs remain largely unclear, primarily due to the absence of this cytokine in rodent models (*8*).

Interestingly, the human IL-26 protein has been demonstrated to exhibit receptor-independent functions. For example, it has been shown that IL-26 can form complexes with DNA and activate intracellular TLR9 in dendritic cells. Furthermore, *in vitro* studies have shown that IL-26 can directly kill bacteria by pore formation in a dose-dependent manner (*9*, *10*), including both pathogenic and commensal strains. Since IL-26 has a direct anti-microbial effect, it could play a role in regulating microbiota composition *in vivo*, with the loss of this function potentially impairing microbiota-dependent gut epithelial homeostasis. Zebrafish possess a single orthologue of the human *IL26* gene (*11*), along with its two receptor chains (*12*). Therefore, this model may provide a suitable system for dissecting the receptor-dependent and -independent functions of IL-26 *in vivo*, including its role in modulating the microbiota and epithelial homeostasis in early life.

In the human gut, IL-26 is produced by lymphocytes, including innate lymphoid cells (ILCs) (*13–16*). ILCs were found to populate the human intestine early in embryonic development, prior to 12 weeks post-conception (*17*), thus preceding gut colonization by the microbiota. We previously demonstrated that ILCs are present in the adult zebrafish gut, displaying a cell type diversity resembling that of human ILCs (*18*). However, whether ILCs are present in the zebrafish gut during early life, whether they respond to the colonizing microbiota by producing IL-26, and the *in vivo* consequences of this expression on gut microbiota and epithelial homeostasis remain unknown.

In this study, we combined zebrafish genetics, transcriptomics, and microscopy with microbiota profiling, gnotobiotics (engineering microbiota composition), and gut bacterial infection tools to uncover a novel role for IL-26 in regulating gut homeostasis during early life. We report that the loss of IL-26 resulted in increased proliferation in gut epithelial progenitors and elevated DNA damage in absorptive enterocytes in the posterior larval gut, a segment that functionally resembles the human colon (*19*). By characterizing these phenotypes in zebrafish lacking the IL-26 receptor and zebrafish reared germ-free, we observed that IL-26 regulates epithelial cell proliferation and DNA damage independently of its receptor-mediated signaling but in a microbiota-dependent manner. Microbiota profiling, combined with microbiota transfer experiments, revealed that epithelial dysregulation in *il26^-/-^* larvae results from gut dysbiosis. Notably, we showed that IL-26 bactericidal activity is conserved in zebrafish and may directly control microbiota composition in zebrafish larvae. Furthermore, IL-26 protected the gut from bacterial infection by regulating bacterial loads, DNA damage, and the immune response. Finally, we identified ILCs as the main cell source of IL-26 in the developing larval gut. Overall, our findings suggest a circuit in which early microbial colonization of the developing gut induces IL-26 production in ILCs, which helps establish a healthy microbiota composition through its intrinsic antibacterial properties, thus maintaining epithelial homeostasis.

## RESULTS

### IL-26 regulates cell proliferation and DNA damage in the zebrafish larval gut

To determine the function of IL-26 in gut homeostasis, we generated *il26*-deficient zebrafish using the CRISPR/Cas9 system (Fig. S1, A and B). *il26^-/-^*fish did not show gross morphological defects (Fig. S1, C and D) and survived to adulthood (3 months post-fertilization) in proportions consistent with Mendelian genetics (Chi-Square = 1.783, *P* value = 0.4101, Data S1).

To characterize IL-26 functions in the developing gut, we took an unbiased approach and performed bulk RNA-sequencing (RNA-seq) on dissected guts from 5 days post-fertilization (dpf) *il26^-/-^*and wild-type (WT) larvae (Fig. 1A). 291 genes were upregulated and 275 downregulated in the guts of *il26^-/-^* larvae (Data S2). To complement our analysis of *il26^-/-^* mutants, we performed transcriptomic analysis upon IL-26 overexpression. To this end, recombinant zebrafish IL-26 (rzIl26) or bovine serum albumin (BSA) were injected into the gut and swim bladder of 5 dpf WT larvae, followed by RNA-seq on dissected guts 1 hour post-injection (hpi) (Fig. 1A). In this condition, 1759 genes were upregulated and 2022 were downregulated (Data S3).

**Fig. 1.**
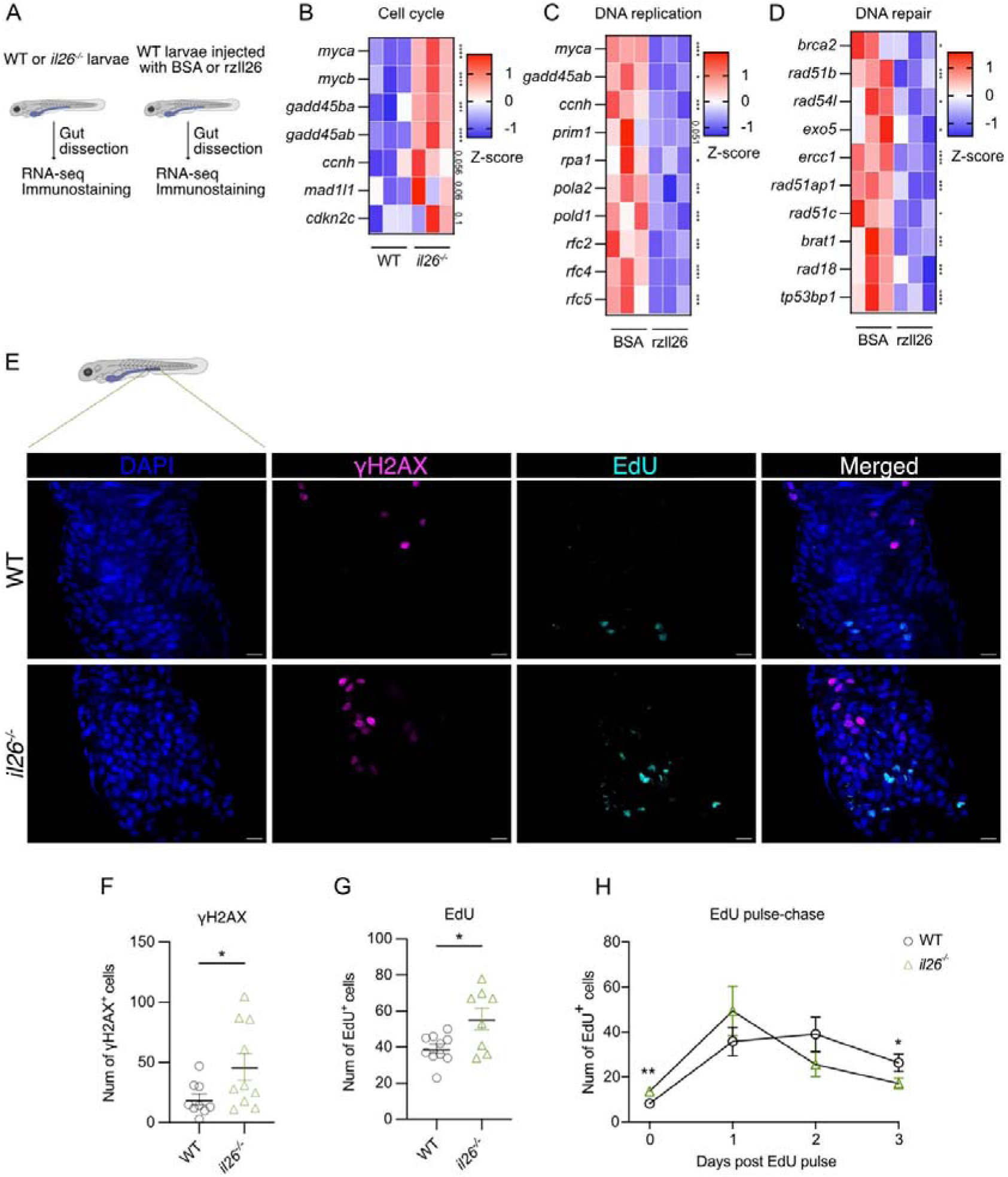
Increased proliferation and DNA damage in the gut of *il26^-/-^* zebrafish larvae. (**A**) A schematic of the experimental setup. Heatmaps of z-scores of genes associated with cell cycle in the loss-of-function dataset (**B**), DNA replication in the overexpression dataset (**C**), and DNA repair in the overexpression dataset (**D**). *P* values are indicated to the right of the heatmaps. (**E**) Representative images of EdU and γH2AX staining in WT and *il26^-/-^*5 dpf posterior larval guts. Scale bars, 10 μm. Quantification of γH2AX (**F**) and EdU (**G**) staining in WT and *il26^-/-^* 5 dpf posterior larval guts. (**H**) EdU pulse-chase analyses in the posterior gut of WT and *il26^-/-^* larvae. Error bars show means ± SEM. Statistical significance was determined by edgeR package in R (B-D), Wilcoxon test (F-H). \**P* < 0.05, \*\**P* < 0.01, \*\*\**P* < 0.001, and \*\*\*\**P* < 0.0001.

To infer the affected biological pathways and processes, we performed Kyoto encyclopedia of genes and genomes (KEGG) and gene ontology (GO) analyses on both datasets (Fig. S1, E-H). KEGG pathway analysis on *il26^-/-^* guts revealed activation of “cell cycle” (Fig. S1E), with upregulation of genes such as *myca* (Myc proto-oncogene a), a transcription factor that promotes proliferation (*20*); and *ccnh* (cyclin H), which controls cell cycle progression and promotes cancer growth (*21–23*) (Fig. 1B). In contrast, KEGG pathway analysis upon rzIl26 injection showed suppression of “DNA replication” (Fig. S1G), with downregulation of genes such as *myca* and *ccnh* (Fig. 1C). Furthermore, GO analysis showed the suppression of “DNA repair” upon rzIl26 injection (Fig. S1H), with downregulation of several DNA repair genes such as *brca2* (breast cancer gene 2 DNA repair associated (*24*)); *rad51b* (RAD51 paralog B (*25*)); and *ercc1* (excision repair cross-complementation group 1 (*26*)) (Fig. 1D). Collectively, our transcriptomic analysis upon IL-26 loss-of-function and overexpression suggested a potential role for IL-26 in regulating cell proliferation and DNA repair in the gut.

To validate these findings, 5 dpf *il26^-/-^* and WT larvae were incubated with EdU for 3 hours, followed by EdU and γH2AX staining to assess proliferation and DNA damage, respectively (Fig. S2A). γH2AX-positive cells were mainly detected in the posterior part of *il26^-/-^* guts. This section of the larval gut is functionally equivalent to the mammalian colon (*19*), the part of the intestine where ulcerative colitis (UC) predominately takes place. Since polymorphisms in the *IL26* gene are a risk factor for UC (*5*), we focused our subsequent analyses on the posterior larval gut. *il26^-/-^* larvae showed a higher number of both γH2AX-positive and EdU-positive cells compared to WT controls (Fig. 1, E-G), indicating elevated DNA damage and cell proliferation in the posterior gut in the absence of IL-26 at this developmental stage.

To better understand the physiological consequences of the observed phenotypes, epithelial turnover was quantified by performing EdU pulse-chase in 5 dpf *il26^-/-^ TgBAC(cldn15la- GFP)* larvae, which specifically labels the basolateral membrane of gut epithelial cells (*27*) (Fig. S2B). In accordance with the results above, *il26^-/-^* larvae showed a higher number of EdU-positive cells than WT at 0 days post-EdU pulse. However, at 3 days post-EdU pulse, the mutants had a lower number of EdU-positive cells compared to WT (Fig. 1H), indicating faster epithelial turnover in the *il26^-/-^* background. Additionally, no differences were observed in the gut lengths of 5 dpf *il26^-/-^* larvae compared to WT (Fig. S2, C and D). Together, our data demonstrate that IL-26 loss leads to increased epithelial cell proliferation, turnover, and DNA damage in the posterior larval gut without causing gross developmental impairments.

### IL-26 suppresses proliferation in epithelial progenitors and DNA damage in absorptive enterocytes

To gain deeper insight into the phenotypes of increased proliferation and DNA damage in *il26^-/-^* guts and to pinpoint the affected cell types, we performed single-cell RNA sequencing (scRNA-seq) on whole dissected guts from 5 dpf *il26^-/-^* and WT larvae. We profiled the gene expression of 13,256 individual cells (7,420 WT; 5,836 *il26^-/-^*) (Fig. 2A), with a median of 1,517 detected genes per cell. We next applied graph-based clustering on our integrated dataset (Fig. 2B) and identified 24 distinct clusters based on the expression of known markers (Data S4).

**Fig. 2.**
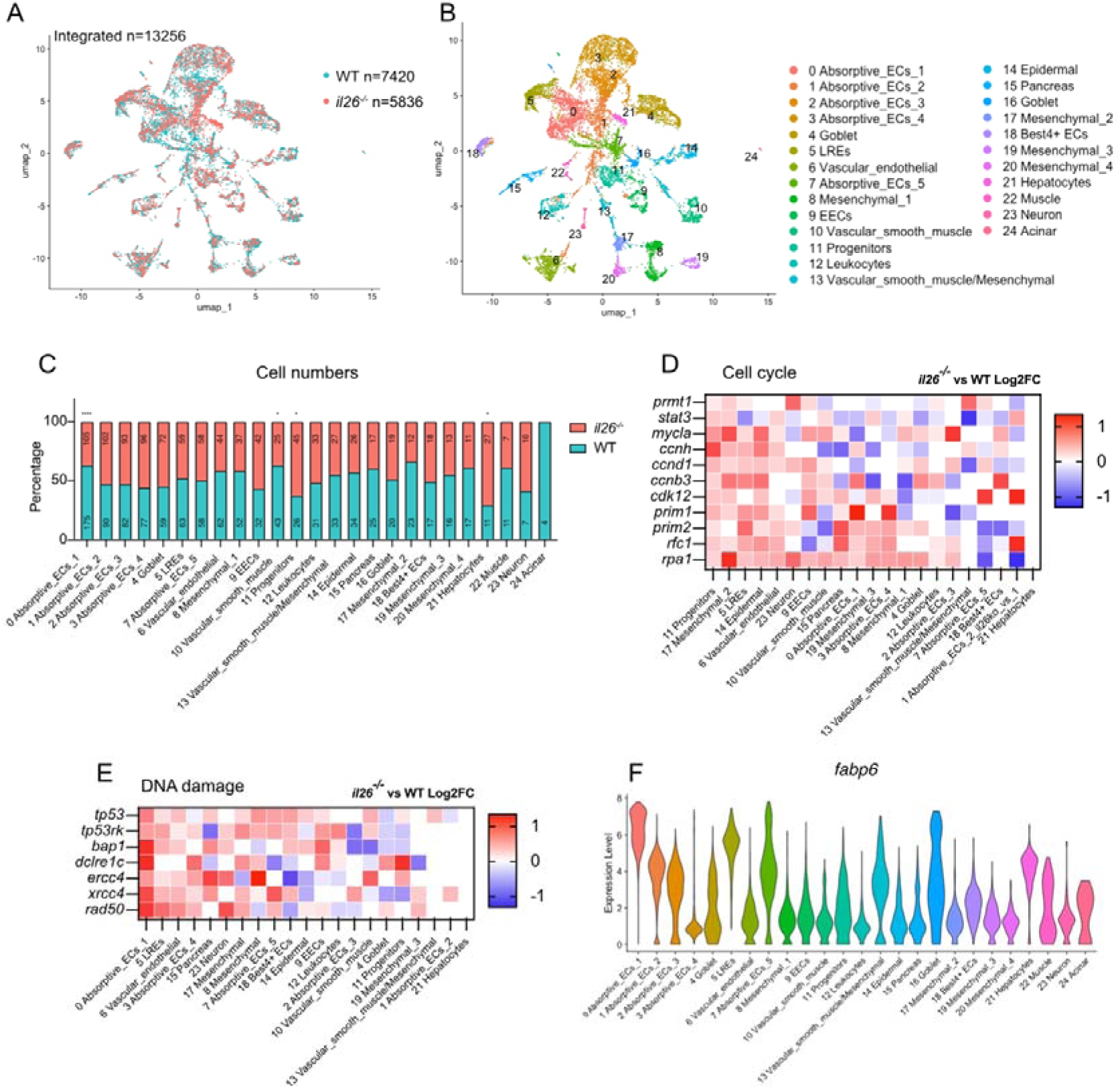
Single-cell RNA sequencing reveals increased proliferation in epithelial progenitors, elevated DNA damage in enterocytes, and altered epithelial cell populations in *il26^-/-^* larval guts. UMAP Dimensional reduction projection of scRNA-seq dataset of dissected guts of 5-dpf *il26^-/-^*and WT larvae. Cells are colored by genotype (**A**) or cell type (**B**). (**C**) Numbers and percentages of detected cells per cell population, normalized to 1000 detected cells. *P* values are indicated when significant. Expression of genes associated with cell cycle (**D**) or DNA damage (**E**), colored by log2FC expression in *il26^-/-^* compared to WT across each cell cluster. (**F**) Violin plot of *fabp6* expression. \**P* < 0.05, \*\**P* < 0.01, \*\*\**P* < 0.001, and \*\*\*\**P* < 0.0001.

In agreement with our cell proliferation observations, the number of epithelial progenitors in *il26^-/-^* guts was higher than in WT (45 vs. 26 per 1,000 detected cells, *P* value = 0.0319) (Fig. 2C). Moreover, epithelial progenitors showed higher expression levels of stemness markers, such as *stat3* (*28*) and *prmt1* (*29*) in *il26^-/-^* compared to WT (Fig. 2D). These findings further reinforce the conclusion that IL-26 loss leads to increased proliferation in epithelial progenitors of larval guts.

Next, to identify the cell types exhibiting higher DNA damage in the guts of *il26^-/-^*, we analyzed expression levels of DNA damage-associated genes such as *tp53* and *rad50* across each cell cluster (Fig. 2E). A cluster of absorptive enterocytes, EC1, which expressed high levels of the posterior enterocyte marker *fabp6* (*19*) (Fig. 2F), displayed heightened expression of these genes (Fig. 2E). Notably, EC1 cells were less abundant in *il26^-/-^* compared to WT (105 vs. 175 per 1,000 detected cells, *P* value < 0.0001) (Fig. 2C). Our *in silico* analysis suggests that absorptive enterocytes of the posterior larval gut accumulate DNA damage in *il26^-/-^*.

To validate these findings, γH2AX staining was performed in *il26^-/-^ TgBAC(cldn15la-GFP)* larvae (Fig. S3A). γH2AX-positive cells were GFP-positive, confirming that DNA damage occurs in gut epithelial cells. Furthermore, co-staining for γH2AX and 2F11, a secretory cell-specific marker in zebrafish (*30*), revealed no colocalization (Fig. S3B). Moreover, *anxa4* (annexin A4), encoding the 2F11 antigen (*31*), showed minimal expression in EC1 cells (Fig. S3C). These results indicate that the loss of IL-26 results in an increase in DNA damage in posterior gut epithelial cells, likely in absorptive enterocytes rather than secretory cells.

Overall, our single-cell transcriptomic and *in situ* analyses suggest that IL-26 specifically suppresses cell proliferation and DNA damage in gut epithelial progenitors and absorptive enterocytes, respectively.

### IL-26 modulation of epithelial homeostasis in the gut is receptor-independent

The IL-26 receptor complex in humans is composed of two subunits: IL-10RB and IL-20RA (*32*, *33*). IL-10RB is shared among IL-10, IL-22, IL-28A, IL-28B, IL-29, and IL-26, whereas IL-20RA is utilized by IL-26, IL-19, IL-20, and IL-24. Notably, the combination of IL-10RB and IL-20RA is unique to IL-26. To elucidate the role of IL-26 receptor signaling in regulating proliferation and DNA damage in gut epithelial cells, we first analyzed the expression profile of these two subunits in our scRNA-seq dataset. Their expression was not detected in either EC1 or epithelial progenitors (Fig. S4, A and B). However, both were expressed in other cell types in the gut, such as EC4 and lysosome rich enterocytes (LREs). To check whether this IL-26 receptor expression plays any role in the phenotypes observed in *il26^-/-^*, we generated *il20ra-*deficient zebrafish using the CRISPR/Cas9 system (Fig. S4, C and D). Interestingly, *il20ra^-/-^* larvae showed similar levels of EdU (Fig. S4, E and G) and γH2AX (Fig. S4, F and G) compared to WT, indicating that IL-26 receptor signaling does not regulate proliferation and DNA damage in the zebrafish larval gut. These findings support the hypothesis that IL-26 regulates epithelial cell proliferation in the gut in a receptor-independent manner.

### Analysis of IL-26 receptor-independent functions reveals conservation of IL-26 intrinsic bactericidal activity in zebrafish

Human IL-26 protein has receptor-independent functions. For example, human IL-26 protein (IL26) can form complexes with DNA and activate intracellular TLR9 in the absence of the IL-26 receptor (*9*). Since TLR9 regulates cell proliferation (*34*, *35*) and DNA damage (*36*), we investigated whether the zebrafish IL-26 protein (Il26) protein can form DNA complexes using gel migration assays (Fig. S5A). In contrast to human IL26, incubating zebrafish Il26 with genomic DNA did not block DNA migration, suggesting that the zebrafish protein does not form complexes with DNA and does not regulate cell proliferation and DNA damage through this property.

However, human IL26 has another receptor-independent function: its intrinsic bactericidal activity. Indeed, human IL-26 has been shown to kill *E. coli* and *P. aeruginosa* but not *E. faecalis*, at a concentration of 8 µM (*9*). We compared the bactericidal activity of human and zebrafish IL-26 proteins by incubating each protein at a concentration of 8 µM with these bacteria, followed by colony-forming units (CFU) analysis. Both proteins killed *E. coli* and *P. aeruginosa* to similar levels but neither protein killed *E. faecalis* (Fig. S5B). These results show that the intrinsic bactericidal activity of IL-26 is conserved in zebrafish, with similar specificity and efficacy.

### IL-26 maintains gut epithelial homeostasis by controlling the composition of the microbiota

Having found that IL-26 regulates gut epithelial cell proliferation and DNA damage in a receptor-independent manner and that zebrafish Il26 protein has bactericidal properties, we wondered whether the lack of this function in *il26^-/-^* might result in an altered microbiota composition, leading to increased proliferation and DNA damage. To determine the role of the microbiota in the observed phenotypes, we first analyzed germ-free (GF) *il26^-/-^* larvae. We performed bulk RNA-seq analysis on dissected guts from 5 dpf WT and *il26^-/-^* larvae reared under GF conditions. 319 genes were upregulated and 307 downregulated in the guts of *il26^-/-^* GF larvae (Data S5). Notably, genes related to cell cycle and DNA repair were not differentially expressed in *il26^-/-^*GF compared to WT GF larvae (Fig. 3A), suggesting that the increased proliferation and DNA damage in *il26^-/-^* is reverted in GF conditions. To validate this, we reared WT and *il26^-/-^* larvae under conventional (CV) or GF conditions and quantified gut cell proliferation and DNA damage. EdU- and γH2AX-positive cells were less abundant in *il26^-/-^* GF compared to *il26^-/-^* CV (Fig. 3B-C and Fig. S6A-B). Furthermore, *il26^-/-^*GF showed similar levels of proliferation and DNA damage as WT GF. These data indicate that IL-26 suppresses cell proliferation and DNA damage in the zebrafish gut in a microbiota-dependent manner.

**Fig. 3.**
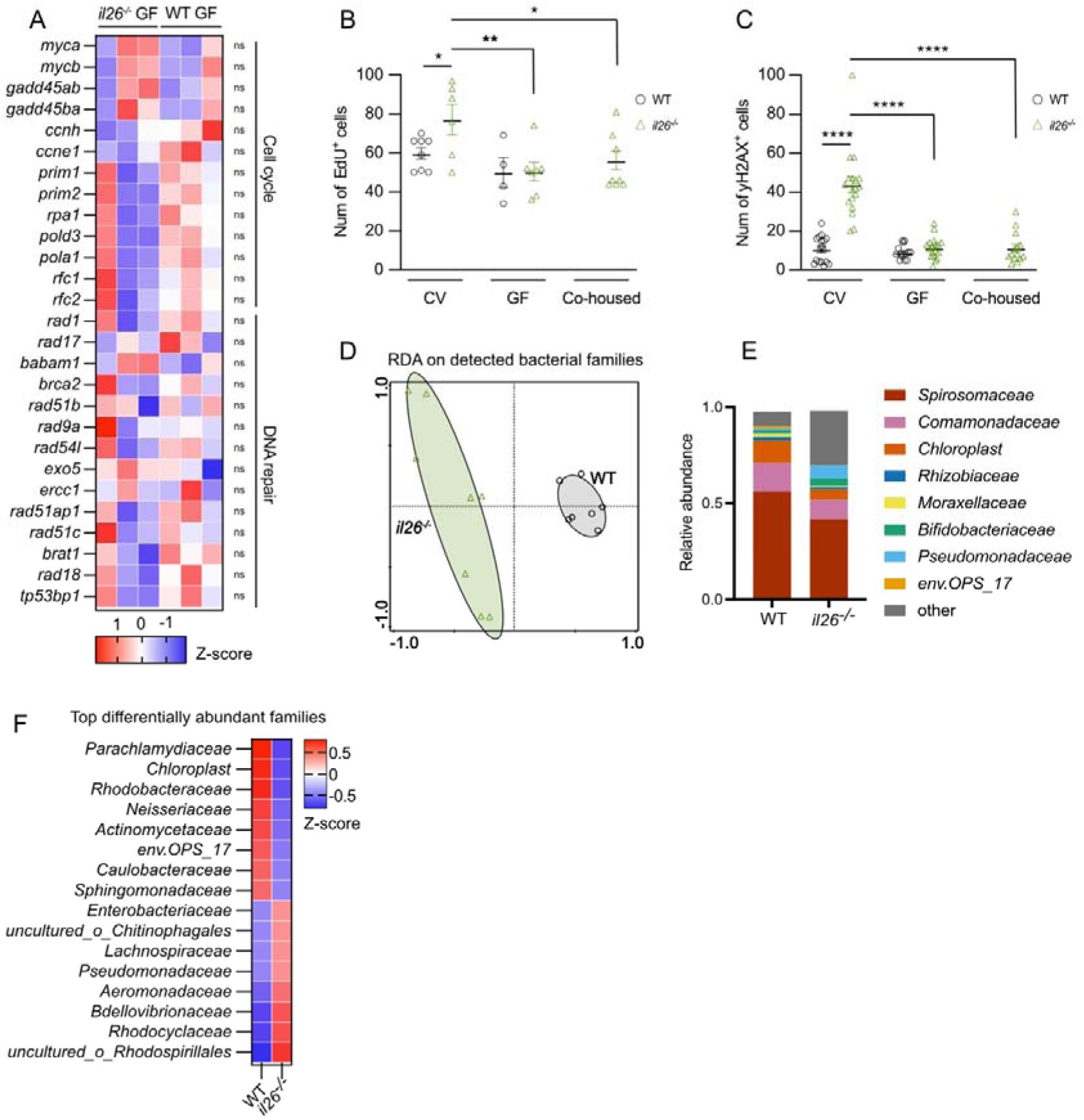
IL-26 regulates epithelial homeostasis by modulating microbiota composition. (**A**) Heatmap of z-scores of genes associated with cell cycle and DNA repair in *il26^-/-^*GF compared to WT GF. *P* values are indicated. Quantification of EdU staining (**B**) and γH2AX staining (**C**) in WT and *il26^-/-^* larval guts reared CV, GF, or cohoused. (**D**) RDA plot on the bacterial families detected by 16S rRNA-sequencing on 5 dpf WT and *il26^-/-^* larval intestines. (**E**) Relative abundance of the detected bacterial families in WT and *il26^-/-^* larval guts. Only families with an abundance greater than 1% in WT are shown. (**F**) Heatmap of z-scores of the relative abundance of the top differentially abundant families in WT and *il26^-/-^*larval guts. Error bars show means ± SEM. Statistical significance was determined by edgeR package in R (A) and 2way ANOVA (B and C). \**P* < 0.05, \*\**P* < 0.01, \*\*\**P* < 0.001, and \*\*\*\**P* < 0.0001.

To uncover whether IL-26 loss leads to dysbiosis in zebrafish larval guts, we profiled the composition of the microbiota using 16S rRNA-seq on dissected guts from 5 dpf WT and *il26^-/-^* larvae. Redundancy analysis (RDA) of the detected bacterial families revealed clear separation between WT and *il26^-/-^* (*P* value = 0.002, permutation test; Fig. 3D, Data S6), pointing to differences in their gut bacterial communities. Indeed, several bacterial families were differentially represented in the WT compared to *il26^-/-^* gut (Fig. 3, E and F). Notably, *Enterobacteriaceae*, correlated with IBD in humans (*37*), were more abundant in *il26^-/-^* larval guts (Fig. 3F). These data demonstrate that IL-26 shapes the composition of the gut microbiota in the zebrafish larval gut.

Next, we reasoned that if the altered microbiota in *il26^-/-^*larvae was responsible for the observed elevated proliferation and DNA damage, then introducing a WT microbiota to *il26^-/-^*larvae might rescue these phenotypes. To verify this, *il26^-/-^*GF larvae were co-housed with WT CV larvae to promote microbiota transfer from WT to mutant animals. This was followed by EdU (Fig. 3B and Fig. S6A) and γH2AX (Fig. 3C and Fig. S6B) staining. Co-housed *il26^-/-^* larvae displayed cell proliferation and DNA damage levels similar to those in WT CV, suggesting that WT microbiota mitigated the increased proliferation and DNA damage in *il26^-/-^*larvae. These observations indicate that *il26*-deficiency leads to dysbiosis, consequently resulting in impaired epithelial homeostasis.

### Innate sensing of the microbiota induces IL-26 expression in the larval gut

Microbial gut colonization is known to induce transcriptional changes, including cytokine expression (*38*, *39*). To determine whether the microbiota influences *il26* expression in the larval gut, mRNA levels were quantified in 5 dpf CV and GF larval guts. GF larval guts showed lower *il26* expression levels compared to WT (Fig. 4A), indicating that the microbiota induces baseline *il26* expression. To uncover the signaling pathways involved in this process, we measured *il26* in larvae deficient for Myd88, a key adaptor protein in TLR signaling (*40*). *il26* expression was lower in *myd88^-/-^* larval guts compared to WT controls (Fig. 4B). These results suggest that microbial colonization in the gut induces baseline *il26* expression in the zebrafish larval gut through TLR-dependent innate immune sensing.

**Fig. 4.**
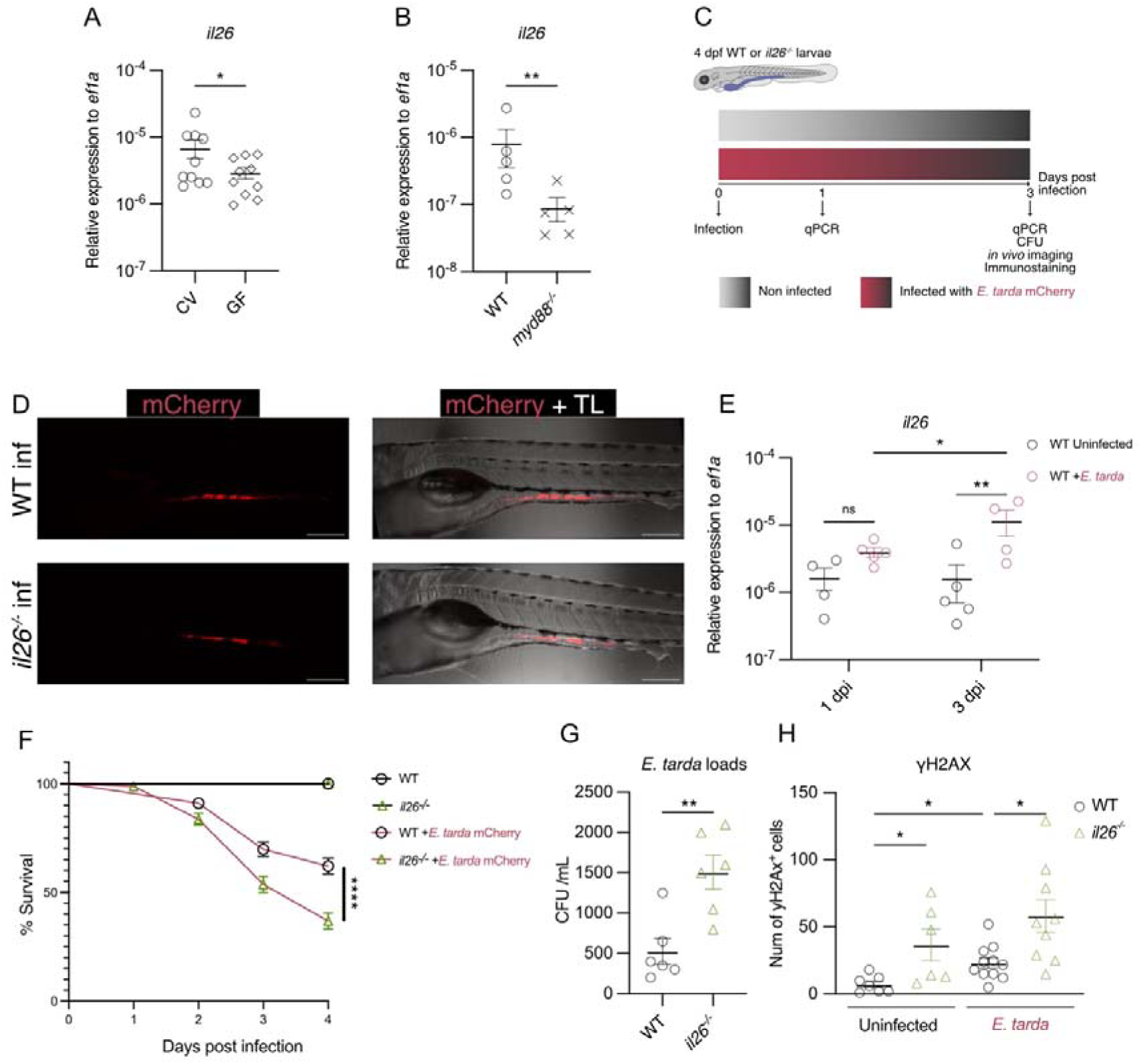
IL-26 expression in the gut is regulated by the microbiota, and *il26^-/-^*larvae are more susceptible to gut bacterial infection. (**A**) qRT-PCR analysis of *il26* in dissected guts of 5 dpf WT larvae reared CV or GF. (**B**) qRT-PCR analysis of *il26* in dissected guts of WT or *myd88^-/-^* 5 dpf larvae. (**C**) Schematic representation of *E. tarda* infection and subsequent analyses. (**D**) WT and *il26^-/-^*larvae infected with mCherry-labeled *E. tarda* at 3 dpi. Scale bars, 200 μm. (**E**) qRT-PCR analysis of *il26* in dissected guts of WT larvae at 1 and 3 dpi. (**F**) Survival analysis of WT and *il26^-/-^* larvae infected with *E. tarda*. (**G**) CFU analysis in dissected guts of WT and *il26^-/-^* larvae at 3 dpi. (**H**) Quantification of γH2AX staining in WT and *il26^-/-^* larval guts at 3 dpi. Error bars show means ± SEM. Statistical significance was determined by Wilcoxon test (A and B), one-way ANOVA (E and H), and Gehan-Breslow-Wilcoxon test (D). \**P* < 0.05, \*\**P* < 0.01, \*\*\**P* < 0.001, and \*\*\*\**P* < 0.0001.

### IL-26 protects the gut from bacterial infection

Since we demonstrated above that zebrafish IL-26 protein possesses intrinsic bactericidal properties, and given that bacterial infections in the gut are known to cause damage and contribute to IBD, we examined whether IL-26 plays a role during gut bacterial infections. We infected WT and *il26^-/-^* larvae via water bath immersion with mCherry-labeled *Edwardsiella tarda*, a Gram-negative bacterium that infects both zebrafish and human guts and is known to cause mortality in zebrafish larvae (*41–43*) (Fig. 4C). Live imaging revealed that *E. tarda* accumulated primarily in the mid and posterior gut at 3 days post-infection (dpi) in both WT and *il26^-/-^* larvae (Fig. 4D). To know whether this infection model induces *il26* expression, we quantified mRNA levels in dissected WT larval guts at 1 and 3 dpi. We found that *il26* expression was higher in 3 dpi larvae compared to uninfected controls (Fig. 4E). Next, we monitored the survival of *E. tarda*-infected WT and *il26^-/-^*larvae. *il26^-/-^* larvae exhibited higher mortality compared to WT upon infection (Fig. 4F). Furthermore, CFU analysis revealed higher bacterial loads in *il26^-/-^* guts at 3 dpi (Fig. 4G). Together, our data show that IL-26 protects the zebrafish developing gut against *E. tarda* infection and mortality.

Next, we investigated the role of IL-26 in regulating gut epithelial proliferation and DNA damage upon *E. tarda* infection by staining WT and *il26^-/-^* larvae with EdU and γH2AX at 3 dpi. *E. tarda*-infected WT and *il26^-/-^* guts showed no differences in the number of EdU-positive cells compared to their respective uninfected controls (Fig. S7, A and B). Interestingly, *E. tarda* infection led to an increased number of γH2AX-positive cells in WT guts (Fig. 4H and Fig. S7A), indicating that *E. tarda* infection induces DNA damage in the posterior larval gut. Moreover, infected *il26^-/-^* guts exhibited greater DNA damage compared to infected WT guts. Our findings indicate that the loss of IL-26 renders gut epithelial cells more susceptible to DNA damage but does not affect their proliferation upon *E. tarda* infection.

Finally, to characterize the immune response in the gut of *il26^-/-^* upon infection, we measured the expression levels of several cytokines in WT and *il26^-/-^* larval guts at 3 dpi. *E. tarda* infection induced the upregulation of *il1b*, *il22*, *tnfa*, and *il10* in WT larvae (Fig. S7, C-F). Similarly, *il26^-/-^* guts showed higher cytokine expression levels upon infection compared to uninfected *il26^-/-^* controls. However, these cytokines showed significantly lower expression levels in *E. tarda*-infected *il26^-/-^* compared to infected WT larvae. In sum, we show that *il26^-/-^*larvae are more susceptible to gut bacterial infection, exhibiting higher bacterial loads, increased DNA damage, and an impaired immune response. These findings underscore the critical role of IL-26 in maintaining gut integrity and regulating immune responses during bacterial infections.

### Innate lymphoid cells are the primary source of IL-26 in the larval gut

Having observed that IL-26–interactions with the microbiota maintain zebrafish gut epithelial homeostasis, we aimed to identify its cellular sources. We first examined *il26* expression in our scRNA-seq dataset, however, it was not detected. This is likely due to the low expression levels of *il26* at steady state as well as the low sequencing depth of current 10X genomics technologies. Since *il26* expression was induced upon inflammation (Fig. 4E), we re- analyzed a published scRNA-seq dataset from guts of larvae incubated with dextran sulfate sodium (DSS), a chemical that induces gut inflammation (*44*). *il26* was mainly expressed in a population of lymphocytes characterized by the expression of *il7r* and *rorc* (Fig. 5, A and B). Interestingly, these cells expressed the novel immune-type receptor 4a gene (*nitr4a*), a specific marker of zebrafish ILCs (*18*) (Fig. 5B). These data suggest that ILCs are present in the larval gut during early life and are the primary source of *il26* expression during inflammation.

**Fig. 5.**
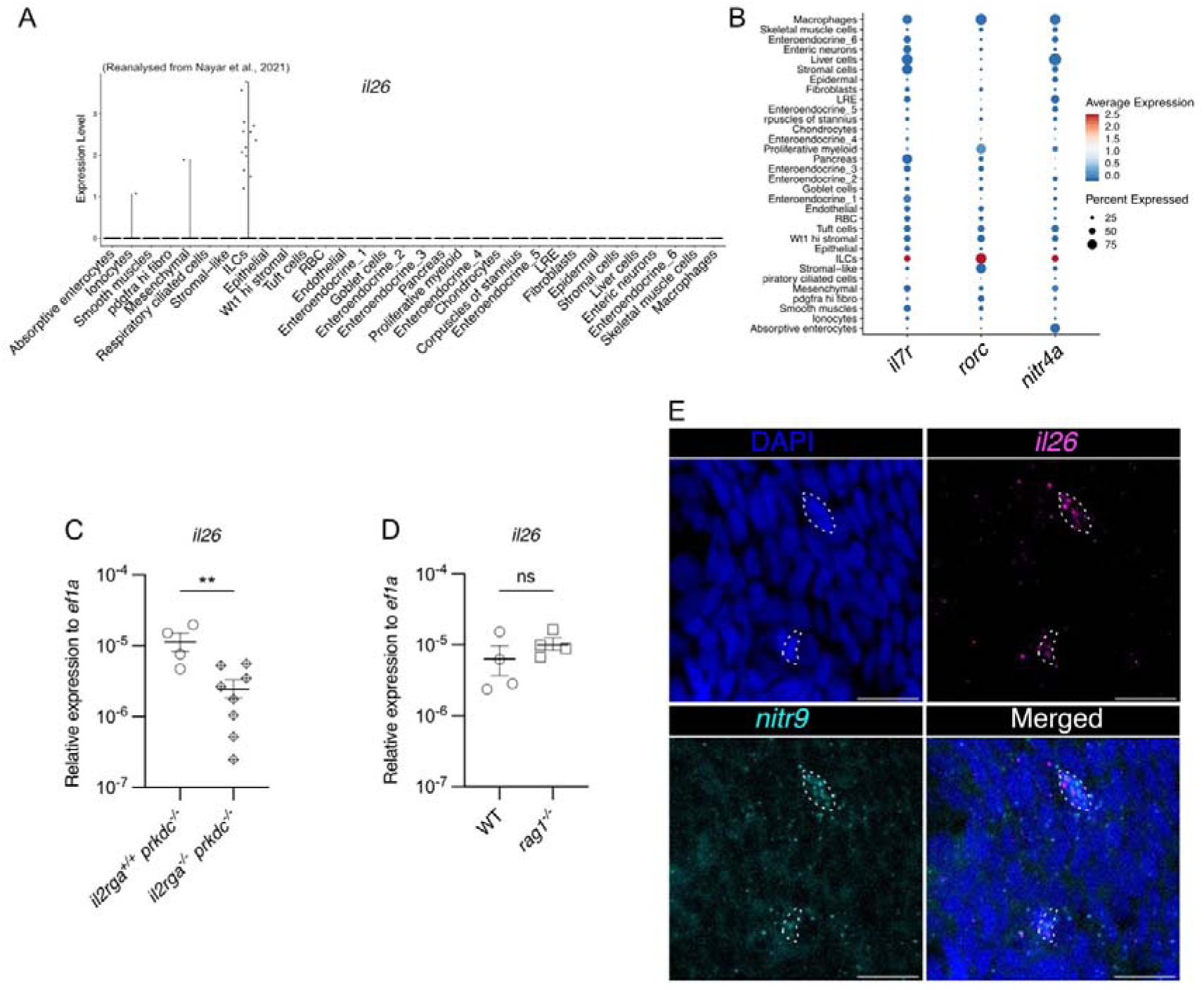
IL-26 is expressed by ILCs in the zebrafish larval gut. (**A**) Violin plot of *il26* expression, reanalyzed from *Nayar* et al., 2021 (***44***). (**B**) Dot plot of ILC markers, reanalyzed from *Nayar* et al., 2021 (***44***). (**C**) qRT-PCR analysis of *il26* in dissected guts of *il2rga^+/+^prkdc^-/-^* and *il2rga^-/-^prkdc^-/-^* 5 dpf larvae. (**D**) qRT-PCR analysis of *il26* in dissected guts of WT or *rag1^-/-^* 5 dpf larvae. (**E**) RNA-FISH of *il26* and *nitr9* on dissected guts of 5 dpf larvae. Scale bars, 10 μm. Error bars in show means ± SEM. \**P* < 0.05, \*\**P* < 0.01, \*\*\**P* < 0.001, and \*\*\*\**P* < 0.0001.

To determine the contribution of ILCs to baseline *il26* expression in the zebrafish larval gut at steady state, complementary approaches were employed. First, we measured *il26* in dissected guts of 5 dpf *il2rga^-/-^prkdc^-/-^* larvae, devoid of adaptive lymphocytes and ILCs (*45*). *il26* levels were lower in *il2rga^-/-^prkdc^-/-^*compared to *il2rga^+/+^prkdc^-/-^* (Fig. 5C), indicating that lymphocytes are required for *il26* expression in the larval gut. Second, we utilized *rag1^-/-^* larvae, which lack adaptive lymphocytes but still possess ILCs (*46*). *il26* levels in *rag1^-/-^* were similar to those in WT larvae (Fig. 5D), indicating that adaptive lymphocytes are not required for *il26* expression. Finally, to visualize *il26* expression *in situ* in gut ILCs, we performed RNA-FISH for *il26* and *nitr9*, a specific marker of zebrafish ILCs (*18*). We detected *il26* expression in *nitr9*-positive cells (Fig. 5E). Together, our data indicate that *il26*- expressing ILCs are present in the developing zebrafish gut and are required for *il26* expression as early as 5 dpf.

## DISCUSSION

In this study, we created the first *in vivo* animal model to study the impact of IL-26 loss-of-function on gut homeostasis. We report that early stages of microbial gut colonization in zebrafish induce IL-26 production in ILCs, which shapes the composition of the microbiota and, in turn, helps maintain epithelial homeostasis (Fig. 6). Our analyses suggest that this function of IL-26 is independent of its canonical receptor complex, composed of IL10Rb and IL20Ra. In addition, we revealed that zebrafish IL-26 possesses intrinsic antibacterial properties, suggesting that the modulation of gut microbiota composition by IL-26 is mediated through this property (Fig. 6).

**Fig. 6.**
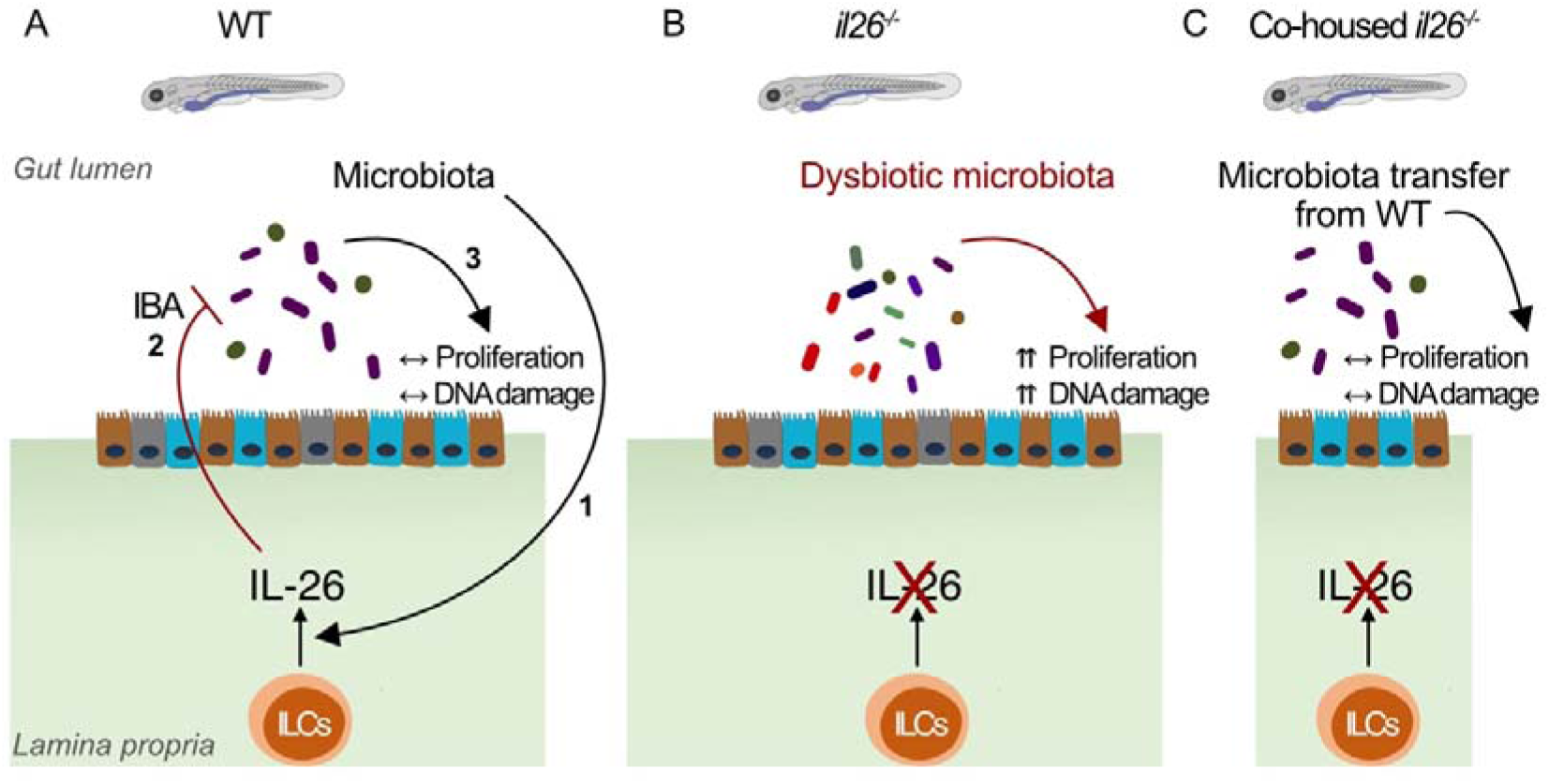
Working model. (**A**) in the developing gut of WT larvae, microbial colonization induces IL-26 expression in ILCs (1), which in turn regulates gut microbiota composition through its intrinsic bactericidal activity (IBA) (2), thereby maintaining baseline levels of proliferation and DNA damage in gut epithelial cells (3). (**B**) In *il26^-/-^* larvae, the loss of IL- 26 IBA leads to dysbiosis, which in turn increases proliferation and DNA damage in the posterior gut epithelium. (**C**) Co-housing mediated microbiota transfer from WT to *il26^-/-^* larvae reascues the high profliferation and DNA damage in *il26^-/-^* gut epithelial cells.

The dysregulation of epithelial homeostasis upon IL-26 loss was marked by heightened cell proliferation in gut epithelial progenitors and elevated DNA damage in posterior absorptive enterocytes, EC1. Notably, our scRNA-seq analysis showed increased numbers of progenitors and decreased numbers of EC1 in *il26^-/-^* (Fig. 2C). These observations underscore how host-microbiota interaction mediated by IL-26 differentially regulates distinct epithelial cell types, paving the way for innovative strategies for targeting the gut epithelium in a cell-specific manner.

Human IL26 is an amphipathic 171-amino acid protein with 30 positively charged residues and an isoelectric point of 10.7. These properties enable receptor-independent functions, such as killing bacteria by pore formation and binding to DNA (*9*). Zebrafish Il26 is an amphipathic 169-amino acid protein with 34 positively charged residues and an isoelectric point of 9.51. The physiochemical properties that the human and zebrafish IL-26 proteins share prompted us to examine the conservation of IL-26 receptor-independent functions in zebrafish. We demonstrated that zebrafish Il26 killed bacteria to similar levels as the human protein, indicating that the intrinsic bactericidal activity of human IL26 is conserved in zebrafish. However, in contrast to human IL26, the zebrafish protein did not bind to DNA. Further analyses of the specific structural and physiochemical properties of human and zebrafish IL-26 proteins may identify the different domains of IL-26 that are involved in receptor binding, DNA binding, and bactericidal activity. This could pave the way for the development of IL-26-derived peptides as novel antibacterial agents, an urgent necessity given the increasing spread of antibiotic-resistant bacteria.

Our 16S rRNA-seq analysis demonstrated that IL-26 deficiency leads to dysbiosis in the zebrafish larval gut. Notably, this was marked by an increased abundance of *Enterobacteriaceae*, a family of bacteria that includes well-known genera such as *Escherichia coli*. This bacterial family is known to colonize the human colon and is consistently enriched in IBD patients (*47*). Moreover, members of the *Enterobacteriaceae* family, such as *pks+ Escherichia coli*, have been reported to produce colibactin, a genotoxin capable of alkylating DNA and inducing double-strand breaks in gut epithelial cells (*48*). Interestingly, we demonstrated that zebrafish IL-26 can kill *E. coli* (Fig. 3D). However, whether the observed increase in DNA damage in the posterior gut of *il26^-/-^* is directly attributable to an enrichment of DNA-damage inducing members of *Enterobacteriaceae* and whether this enrichment is a consequence of the loss of IL-26 antibacterial function warrants further investigation. This may inform the development of IL-26 supplementation as a potential therapeutic strategy for IBD patients with elevated levels of *Enterobacteriaceae*.

IBD patients carrying the *IL26* risk allele exhibit lower serum levels of IL26. Consistently, peripheral blood mononuclear cells (PBMCs) from these patients display a reduced ability to produce IL26 and to kill *E. coli in vitro* (*49*). However, the gut microbiota composition of these specific patients remains uncharacterized. Investigating whether these patients exhibit dysbiosis, particularly with increased levels of *Enterobacteriaceae*, would be of interest. Notably, we observed that epithelial homeostasis in *il26^-/-^* guts can be restored by the transfer of microbiota from WT to *il26^-/-^* larvae, highlighting the therapeutic potential of microbiota transfer in genetically predisposed IBD patients.

The prevalence of IBD has been increasing in the young population (*50*), emphasizing the critical role of a balanced gut ecosystem from early life. Our findings suggest that IL-26 deficiency can influence microbiota composition and disrupt gut homeostasis during early life. These results pave the way for investigating whether early-life IL-26 levels could serve as predictive markers for IBD, offering a significant diagnostic advantage. Furthermore, future research could explore the potential of IL-26 supplementation in at-risk young individuals as a preventive strategy against IBD.

Our findings establish that IL-26 protects the gut from bacterial infections, and that this correlates with increased bacterial loads and DNA damage as well as an impaired immune response. This study underscores the potential of using IL-26 as an agent to fight off bacterial infections in the gut.

ILCs are known to regulate gut homeostasis through cytokine production in the adult zebrafish gut (*16*). However, the emergence of ILCs in the gut during early life and their cytokine production profile were previously unknown. We report that functional ILCs appear in the larval gut as early as 5 dpf as the major source of IL-26. This study underscores the importance of ILCs and the cytokines they produce in gut homeostasis during the first steps of gut microbial colonization in early life.

In summary, our findings reveal key mechanisms by which host-microbiota interactions during early development, mediated by ILC-produced IL-26, protect against dysbiosis, excessive cell proliferation, and DNA damage in epithelial cells.

## MATERIALS AND METHODS

### Zebrafish lines and husbandry

The zebrafish lines: wild-type (AB), *TgBAC(cldn15la-GFP)* (*27*)*, rag1^-/-^* (*51*), *myd88^-/-^* (*52*), *il2rga^-/-^prkdc^-/-^* (*45*), *il26^-/-^,* and *il20ra^-/-^* were reared and kept in the zebrafish core facility at the Institut Curie animal facility in accordance with European Union regulations on laboratory animals using protocol numbers: APAFIS#27495-2020100614519712 v14.2.2, #2019_010 and #2022-008 (approved the French Ministry of Research). Zebrafish larvae used in collaborations with labs in other countries were maintained according to the “Acta de Aprobación 004/2021” provided by the Universidad Andrés Bello, Chile. Zebrafish embryos were collected by natural spawning of adults and were kept at 28°C in E3 water.

### Generation of *il26*- and *il20ra*-deficient zebrafish

The coding sequence for the zebrafish interleukin-26 gene (gene name: *il26*, ENSEMBL ID: ENSDARG00000045672.6) was targeted using CRISPR/Cas9 technology with two specific sgRNAs: GCAGGGATTTATGGATGTCC and GAGACAATAAACCCTTCCAT. Interleukin-20 receptor A (gene name: *il20ra*, ENSEMBL ID: ENSDART00000043626.6) was targeted using two specific sgRNAs: TGGACGTCTCGCGGCTCAGG and GTGAAGTGGACGGCAGGACA. One-cell stage zebrafish embryos were injected with 1 nL of a mixture containing guide RNA (6.65 μM) and Cas9 protein (5 μM).

### Genotyping

Adult zebrafish were anesthetized with tricaine (100 ug/ml, Sigma, #A5040), their tails were cut and incubated for 1 hour at 56°C with FinClip buffer (10 mM Tris, pH 8.0, 10 mM EDTA, 200 mM NaCl, 0.5% SDS) containing Proteinase K (0,2 mg/mL, Invitrogen, #25530- 049). DNA was precipitated by adding 70% ethanol. The pellet was resuspended in water and the product solution was used for genotyping. Zebrafish larvae were anesthetized with tricaine, their tails were cut and incubated during 15 min at 95°C in Base buffer (25 mM KOH, 0.2 mM EDTA). An equal volume of Neutralization buffer (40 mM Tris-HCl) was then added, and the solution was used for genotyping. *il26^-/-^* fish were genotyped by gel electrophoresis using the primers: GTCAAAAGTGAGGTTGTGGCA and CCATGAATGCAGCCTTCAGC. *il20ra^-/-^* fish were genotyped by sequencing using the primers GTTGTGGCTGCTGTACGCTA and GGAACAGGGTTGGGAAGCTAAA.

### Zebrafish Il26 protein injections

Injections were performed using 2 μL of recombinant zebrafish IL-26 protein (1 mg/mL, Kingfisher Biotech, #RP1773Z-025) mixed with 0.5 μL of phenol red. BSA at 1 mg/mL was used as a control.

### Bulk RNA-sequencing

10-15 guts per replicate were dissected and RNA was extracted using the Single Cell RNA Purification Kit (Norgen, #51800) following the manufacturer’s instructions. RNA integrity and concentration were analyzed on the Agilent 4200 Tapestation system using the High Sensitivity RNA ScreenTape Analysis kit (Agilent, #5067-5579). RNA sequencing libraries were prepared from 500 ng to 1 μg of total RNA using the Illumina TruSeq Stranded mRNA Library Preparation Kit. cDNA quality was checked on the Agilent 2100 Bioanalyzer using the Agilent High Sensitivity DNA Kit (Agilent #5067-4626). After quality control, libraries were sequenced with 100-bp paired-end (PE100) reads on the NovaSeq 6000 (Illumina) sequencer. Raw data were checked for quality using FastQC (v0.11.8) and aligned to the reference genome for *Danio rerio* danRer11 from the Genome Reference Consortium. Analysis was performed in R using the EdgeR (*53*) and ClusterProfile (*54*) packages.

### Immunostaining on dissected larval guts

Larvae were incubated with 100 µM EdU in E3 for 3 hours. After incubation, larvae were washed with E3. Next, the guts were dissected and fixed in 4% paraformaldehyde for 1 hour at room temperature. Samples were washed twice with PBST (0.1% Triton-X100 in PBS), followed by incubation in PBS with 3% BSA for 1 hour. After one wash with PBS, the samples were incubated overnight at 4°C with primary antibodies diluted in 200 ul of PBS: γH2AX (1:200, GeneTex, #GTX127342) or 2F11 (1:200, Abcam, #ab71286). The next day, the samples were washed three times for 10 minutes each in PBST. They were then incubated with secondary antibodies diluted in 500 ul of PBS at 4°C for 3 to 4 hours: Goat Anti-Rabbit IgG H&L (1:500, Abcam, #ab175471) or Donkey Anti-Rabbit IgG H&L (1:500, Abcam, #ab150075). Following this, the samples were washed twice for 10 minutes in PBST, and then washed three more times for 10 minutes in PBS. The Click-iT™ EdU Cell Proliferation Kit (Invitrogen, #C10337, # C10340, #C10638) was used according to the manufacturer’s instructions. Samples were washed twice in PBS for 10 minutes. The dissected guts were then mounted with ProLong™ Gold Antifade Mountant (ThermoFisher, #P36931). Images were acquired using the Upright Spinning Disk Confocal Microscope (Roper/Zeiss) with a 63X objective (63x/1.4 OIL DICII PL APO, 420782-9900). Multi-dimensional imaging of the posterior gut was performed to encompass the entire intestinal tube. Quantification was carried out in ImageJ.

### Gut length analysis in larvae

Larvae were anesthetized with tricaine and mounted in 3% methylcellulose for live imaging. Gut length was measured from the intestinal bulb to the end of the intestine at the anal pore. The analysis was done in ImageJ.

### Adult body length measurements

Adult fish were anesthetized with tricaine. Body length was measured from the head to the tail, excluding the tail fin using a ruler.

### Single-cell RNA-sequencing

10 guts from 5 dpf larvae were dissected and placed in 200 μL of a dissociation cocktail (1 mg/mL fresh collagenase A, 40 μg/mL proteinase K, and 0.25% trypsin in PBS) at 37°C for 30 minutes. 50 guts were dissected per condition for a total of 5 replicate tubes per condition. Cells were centrifuged at 300 RCF for 15 minutes at 4°C. The pellet was resuspended in 200 μL PBS with 0.04% non-acetylated BSA. Replicates were pooled and filtered through a 40 μm cell strainer (Fisherbrand, #22363547). Samples were centrifuged at 300 RCF for 15 minutes at 4°C and resuspended with in 50 ul of PBS with 0.04% non-acetylated BSA. To isolate live cells, an OptiPrep™ Density Gradient Medium (Sigma, #D1556-250ML) was used. A 40% (w/v) iodixanol working solution was prepared by mixing 2 volumes of OptiPrep™ with 1 volume of 0.04% BSA in 1X PBS/DEPC-treated water. This working solution was used to create a 22% (w/v) iodixanol solution in the same buffer. The cell suspension was mixed with one volume of the working solution and 0.45 volume of the cell suspension via gentle inversion, transferred to a 15 mL conical tube, and topped up to 6 mL with the working solution. This solution was overlaid with 3 mL of the 22% iodixanol, and then with 0.5 mL of PBS with 0.04% BSA. Samples were centrifuged at 800 RCF for 20 minutes at 20°C. Viable cells were collected from the top interface, diluted in PBS with 0.04% BSA, and centrifuged at 300 RCF for 10 minutes at 4°C. The supernatant was discarded, and the cells were resuspended in PBS with 0.04% non-acetylated BSA to reach the desired concentration. Cells were loaded onto a 10× Chromium instrument (10× Genomics) and libraries were prepared using the Single Cell 3′ Reagent Kit (V2 chemistry) (10× Genomics) according to the manufacturer’s instructions. The sequencing coverage was approximately 100,000 reads per cell. Data analysis was performed using the Seurat package in R (*55*).

### Gel migration assay to determine DNA-binding ability of IL-26

Genomic zebrafish DNA was mixed with different cytokines at a final concentration of 3 ng/µl. Samples were loaded onto a 1.5% agarose gel for electrophoretic migration.

### Antimicrobial assay

The following bacterial strains were used: *Pseudomonas aeruginosa*, *Escherichia coli*, and *Enterococcus faecalis*. Bacteria were cultured at 37°C overnight in trypticase soy broth with 10 mM NaCl. Samples were subcultured for an additional 3 hours to achieve mid–logarithmic phase growth. Bacterial concentrations were measured by spectrophotometry at 620 nm and diluted to a final concentration of 10^5^ CFU/ml, after which they were incubated at 37°C for 24 hours under low-ionic-strength conditions (10 mM NaCl). Human or zebrafish IL-26 were added to these cultures to test their antimicrobial activity. After 24 hours, serial dilutions of bacterial cultures were plated onto lysogeny broth (LB) agar plates overnight. The number of colonies was counted manually.

### Generation of germ-free larvae and co-housing experiments

Fertilized zebrafish eggs were treated with bleach (0.05%) for a maximum of 2 minutes at 3 to 4 hours post fertilization and then washed twice with sterile E3 medium for 5 minutes. Embryos were incubated in chlorine hypochlorite (0.003%) for 20 minutes. After washing, embryos were transferred to sterile E3 medium containing ampicillin (200 μg/mL), kanamycin (5 μg/mL), ceftazidime (200 μg/mL), and chloramphenicol (20 μg/mL) and placed at 28°C in isolated containers. The medium was renewed daily under sterile conditions until the day of sample collection. Sterility was monitored every 2 days by incubating fish water in TBS media for 24 hours at 37°C. Cohousing experiments were performed by transferring GF larvae to containers with CV larvae, separated by a sterile strainer (Fisherbrand, #22363547).

### Microbial DNA extraction and 16S rRNA sequencing

20 guts from WT and *il26^-/-^* 5 dpf larvae per replicate were dissected. Bacterial DNA was isolated using the DNeasy PowerSoil kit (Qiagen, #47014). Amplification and barcoding was performed using V3-V4 primers (V3: CCTACGGGNGGCWGCAG; V4: GACTACHVGGGTATCTAATCC) using the LongAmp Taq DNA polymerase (NEB, M0323S) using the following PCR program: 94[C 30 sec, 30x (94[C 30 sec, 54[C 1 min, 65[C 40 sec), 65[C 10 min, 12[C. After PCR clean-up (Invisorb Fragment CleanUp; Invitek 1020300300) samples were sequenced using Oxford Nanopore (MinION) using the ligation sequencing kit V14 (SQK-LSK114) and R10.4.1 flow cell. The raw sequencing data (pod5 files) were basecalled to fastq files using the super accuracy model of Dorado v0.5.3.(*56*). Reads were demultiplexed and subsequently filtered by size, retaining those with lengths ranging from 300 to 700 base pairs. Demultiplexed reads were analysed with Qiime2 (*57*),using DADA2 for ASV calling and the Silva database version 138 for taxonomic classification. Redundancy analysis was done with Canoco 5.15 (*58*) with bacterial microbiota composition (relative abundance at the family level) as response variables, after transforming with the formula log (1000*relative_abundance[+[1). RDA *P* values were determined through permutation testing (500 permutations).

### Quantitative real-time PCR

10 guts per replicate were dissected and RNA was extracted using the Single Cell RNA Purification Kit (Norgen, #51800) following the manufacturer’s instructions. Synthesis of cDNA was performed using the M-MLV Reverse Transcriptase Kit (Invitrogen, #28025013). qPCR was carried out using the Takyon™ Kit (Takyon, #UF-LPMT-C0701) on a Thermo ABI ViiA 7 Real-Time PCR System (Thermo Applied Biosystems). The primers used were as follows: *il26*: CCATAAATCCCTGCCGAGAGA, CACGCTTGAAGTCTGGGACA; *il20ra*: GGAGTACGCCATTTACGGGG, ACAGTGGTCTGAATCTGCCG; *il1b*: GAGACAGACGGTGCTGTTTA, GTAAGACGGCACTGAATCCA; *tnfa*: GGCCTTTTCTTCAGGTGGCT, AGCACTTGTTCCTCAGTCAGT; *il22*: TGCAGAATCACTGTAAACACGA, CTCCCCGATTGCTTTGTTAC; *il10*: TCACGTCATGAACGAGATCC, CCTCTTGCATTTCACCATATCC.

### *E. tarda* infections

*Edwarsiella tarda* FL60 (provided by Dr. Phillip Klesius (USDA, Agricultural Research Service, Aquatic Animal Health Research Unit) were grown in TSB medium + tetracycline (15 ug/mL) at 28°C overnight. A 1:100 dilution was performed, and the bacteria were grown to reach OD600 = 0.250. The bacteria were then centrifuged at 3500 RCF for 5-10 minutes and resuspended in E3 water to achieve OD600 = 0.250. 6 ml of the bacterial suspension in E3 were added to six larvae for 5 hours at 28°C. Larvae were washed three times with E3 water. Survival was monitored every 12 hours for 3 days post-infection. Images were acquired with the THUNDER Imager Model Organism microscope (Leica). mCherry area and mean intensity were determined using ImageJ.

### Colony-forming units (CFU) analysis

Five guts were dissected per replicate in 100 µL of PBS. The tissue was dissociated by vortexing. 1:10 dilution was prepared, and 20 µL of the diluted sample was spread onto LB-agar plates supplemented with tetracycline (15 µg/mL). Plates were incubated overnight at 37°C, and colonies were counted manually.

### HCR™ RNA-FISH

HCR probes for zebrafish *il26* and *nitr9* were purchased from Molecular Instruments. The staining was performed according to the manufacturer’s guidelines with the following optimizations: guts were dissected and fixed in 4% paraformaldehyde for 1 hour at room temperature. After completing the staining protocol, the dissected guts were mounted with ProLong™ Gold Antifade Mountant (ThermoFisher, #P36931). Images were acquired using the Upright Spinning Disk Confocal Microscope (Roper/Zeiss) with a 63X objective (63x/1.4 OIL DICII PL APO, 420782-9900).

### Statistical analysis

Statistical analyses were performed using RStudio or GraphPad Prism. Specific statistical tests and significance levels are detailed in the respective figure legends.

## Acknowledgments

The authors would like to acknowledge the members of the animal facility of the Institut Curie for zebrafish care. Life Science Editors provided editing services for this manuscript.

## Funding

This work was supported by the Institut Curie, INSERM, and CNRS, and the grants listed below.

Laboratoire d’Excellence (Labex) DEEP (ANR-11-LBX-0044, ANR-10-IDEX- 0001- 02 PSL) (PPH)

Ville de Paris Emergence Program (2020 DAE 78) (PPH) FRM amorçage (AJE201905008718) (PPH)

ATIP-Avenir Starting Grant R21045DS (PPH)

ERC-StG Cytok-Gut 101041422 (PPH, SR, AJK, YS)

PhD fellowships from the Ministère de l’Enseignement Supérieur et de la Recherche and from the Fondation pour la Recherche Médicale (FDT202304016654) (YS)

## Author contributions

Conceptualization: YS, PPH

Methodology and Investigation: YS designed and conducted all experiments with the following contributions: GG assisted with germ-free experiments and qPCRs; KQC and CF assisted with E. tarda infections; SR and CGB assisted with gut dissections; AJK optimized the single-cell suspension protocol for the scRNA-seq experiment; SB and JB performed and analyzed the 16S rRNA sequencing experiments; DPP and JMG conducted the in vitro antimicrobial assays; RMC and EJV contributed to the analysis of the scRNA-seq dataset from *Nayar* et al., 2021.

Visualization: YS

Funding acquisition: YS, PPH

Project administration: YS, PD, GG, PPH Supervision: PPH

Writing – original draft: YS

Writing – review & editing: YS, PPH

## Data and materhials availability

The new reagents and datasets generated during the current study are available from the corresponding author upon request.

