## Supplementary materials for "IL-26 from ILCs regulates early-life gut epithelial homeostasis by shaping microbiota composition"

Yazan Salloum *et al.*

**This file includes:**

Figs. S1 to S7

**Other Supplementary Materials for this manuscript include the following:**

Data S1 to S6

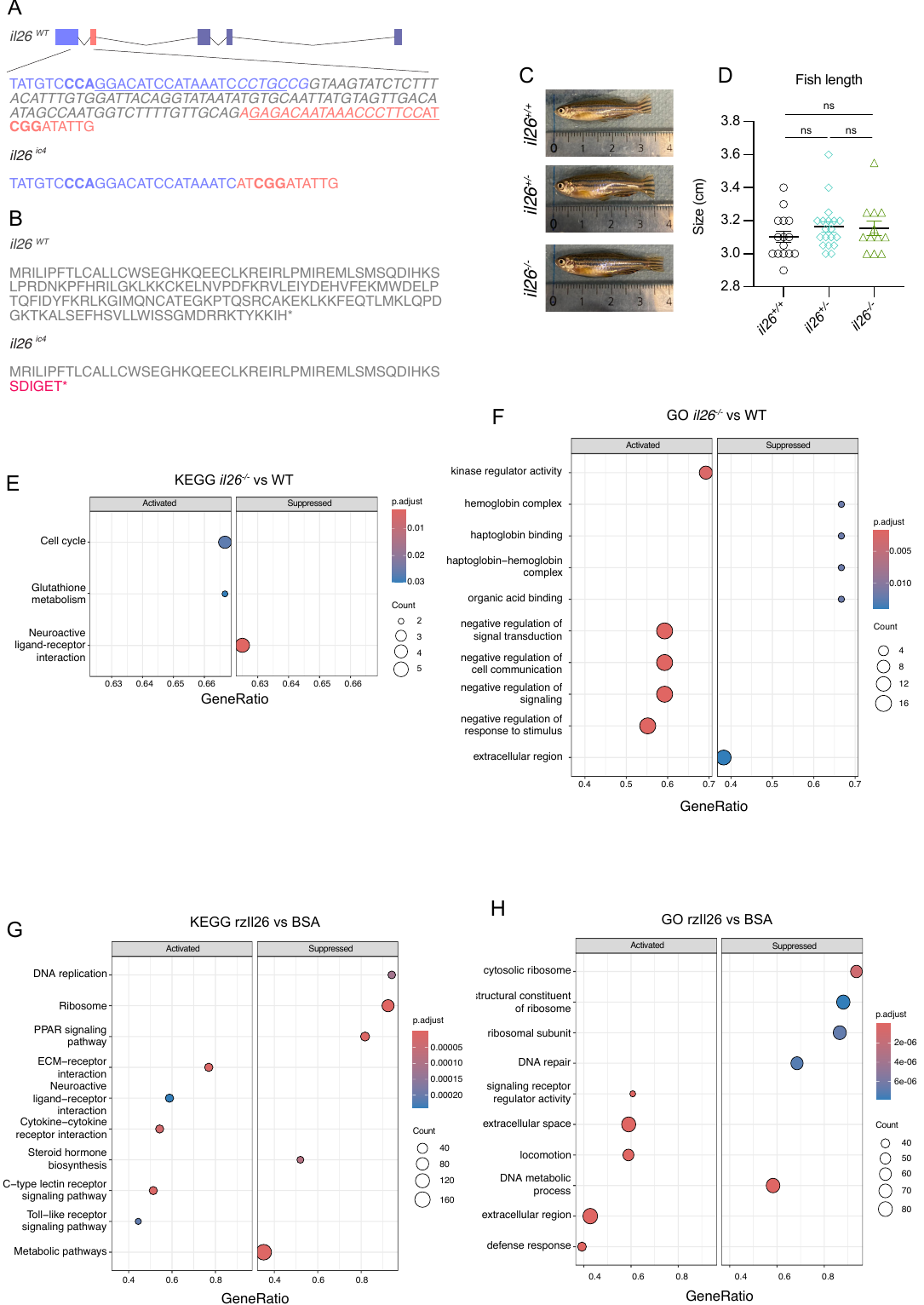

**Fig. S1: Generation and characterization of *il26*-deficient zebrafish and an IL-26 overexpression model.** (**A**) Schematics and DNA sequence of the WT and mutant zebrafish *il26* alleles. The single guide RNA (sgRNA) target sites are underlined. The protospacer adjacent motif (PAM) sequences are indicated in bold. The deleted nucleotides in *il26^ic4^* are highlighted in *italics.* Our approach resulted in a 110-bp deletion across exon 1 and exon 2 of the zebrafish IL-26 gene (*il26^ic4^*). (**B**) The predicted protein sequence of *il26^ic4^* consisting of 53-amino acids. Images of *il26^+/+^*, *il26^+/-^*, and *il26^-/-^* adult fish (**C**) and their length (**D**). KEGG pathway (**E**) and GO analysis (**F**) on the loss-of-function dataset. KEGG pathway (**G**) and GO analysis (**H**) on the overexpression dataset. Error bars show means ± SEM. Statistical significance was determined by Kruskal-Wallis test (D). ns *P* > 0.05.

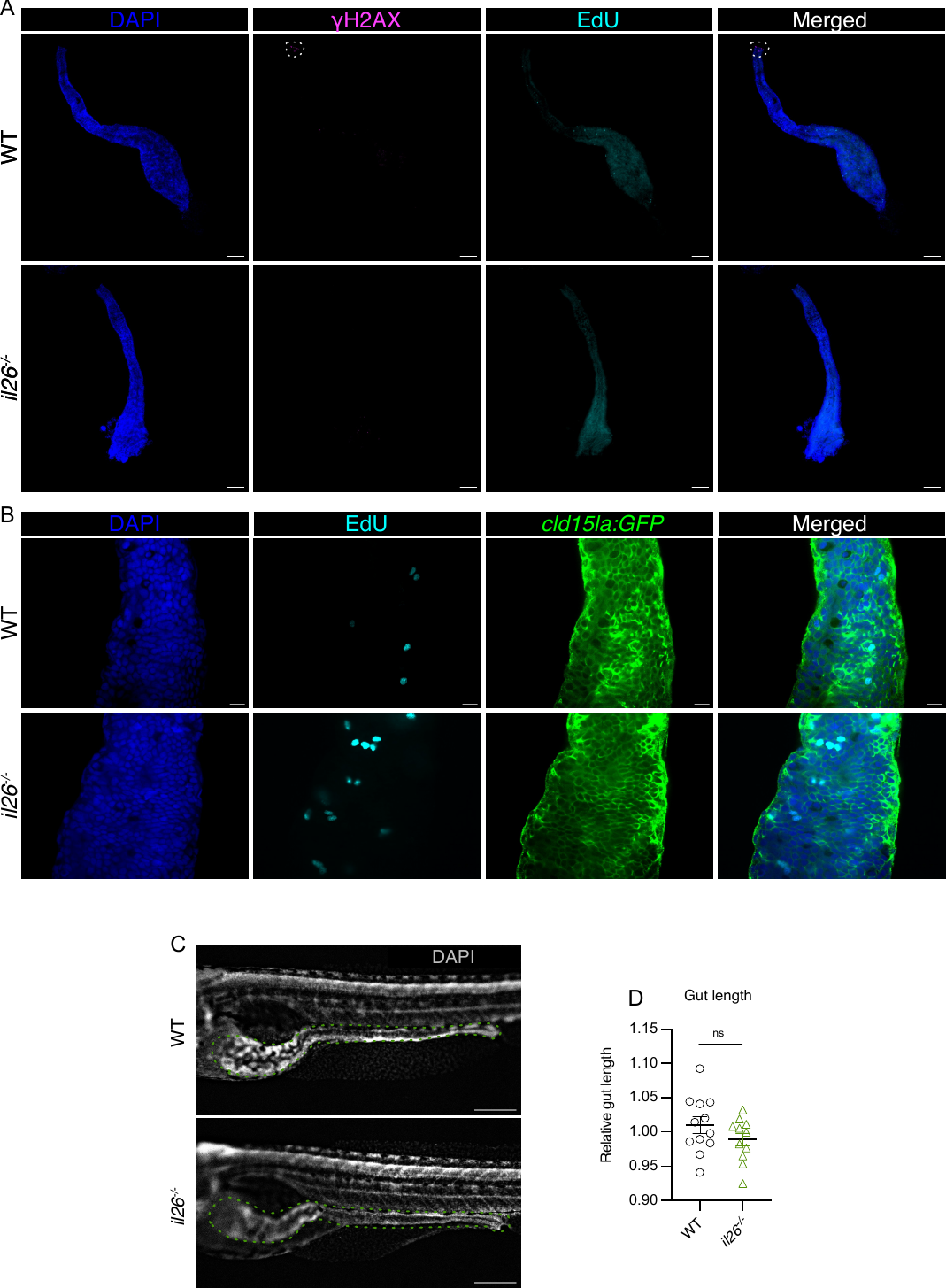

**Fig. S2:** **Immunostaining in *il26^-/-^* larval guts.** (**A**) Immunostaining for EdU and γH2AX of 5 dpf WT and *il26^-/-^* larvae. Scale bars, 100 μm. (**B**) EdU staining of 5 dpf *TgBAC(cldn15la-GFP)* in WT and *il26^-/-^* genetic backgrounds. Scale bars, 10 μm. Images of WT and *il26^-/-^* larvae (**C**) and their gut length (**D**). Scale bars, 100 μm. *^-/-^*. Error bars in show means ± SEM. Statistical significance was determined by Wilcoxon test (D) and Kruskal-Wallis. ns *P* > 0.05.

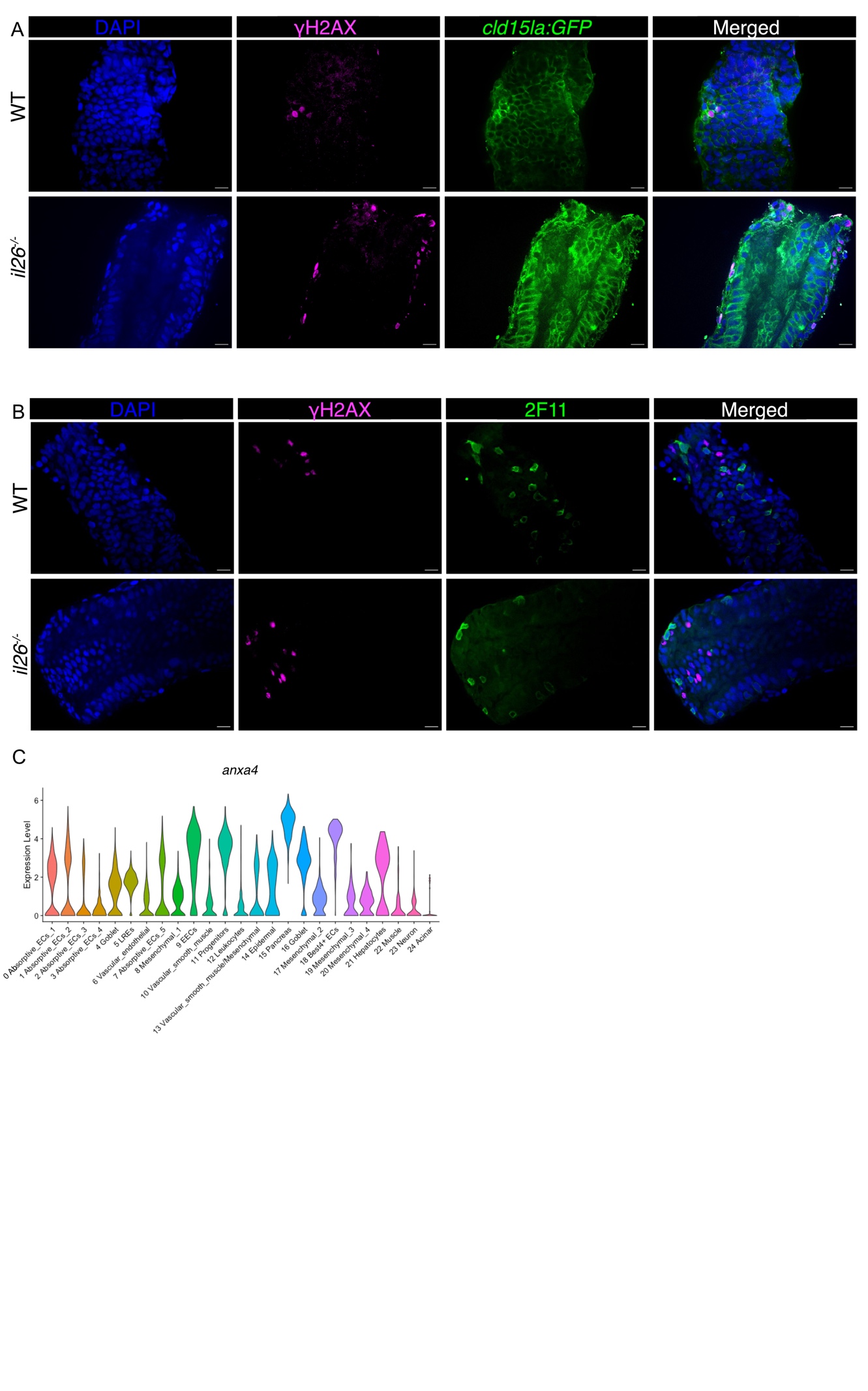

**Fig. S3: Characterization of the higher proliferation and DNA damage observed in the guts of *il26^-/-^* larvae.** (**A**) γH2AX staining of 5 dpf *TgBAC(cldn15la-GFP)* in WT and *il26^-/-^* genetic backgrounds. Scale bars, 10 μm. (**B**) Co-immunostaining for γH2AX and 2F11in WT and *il26^-/-^*. Scale bars, 10 μm. (**C**) Violin plot of *anxa4* expression.

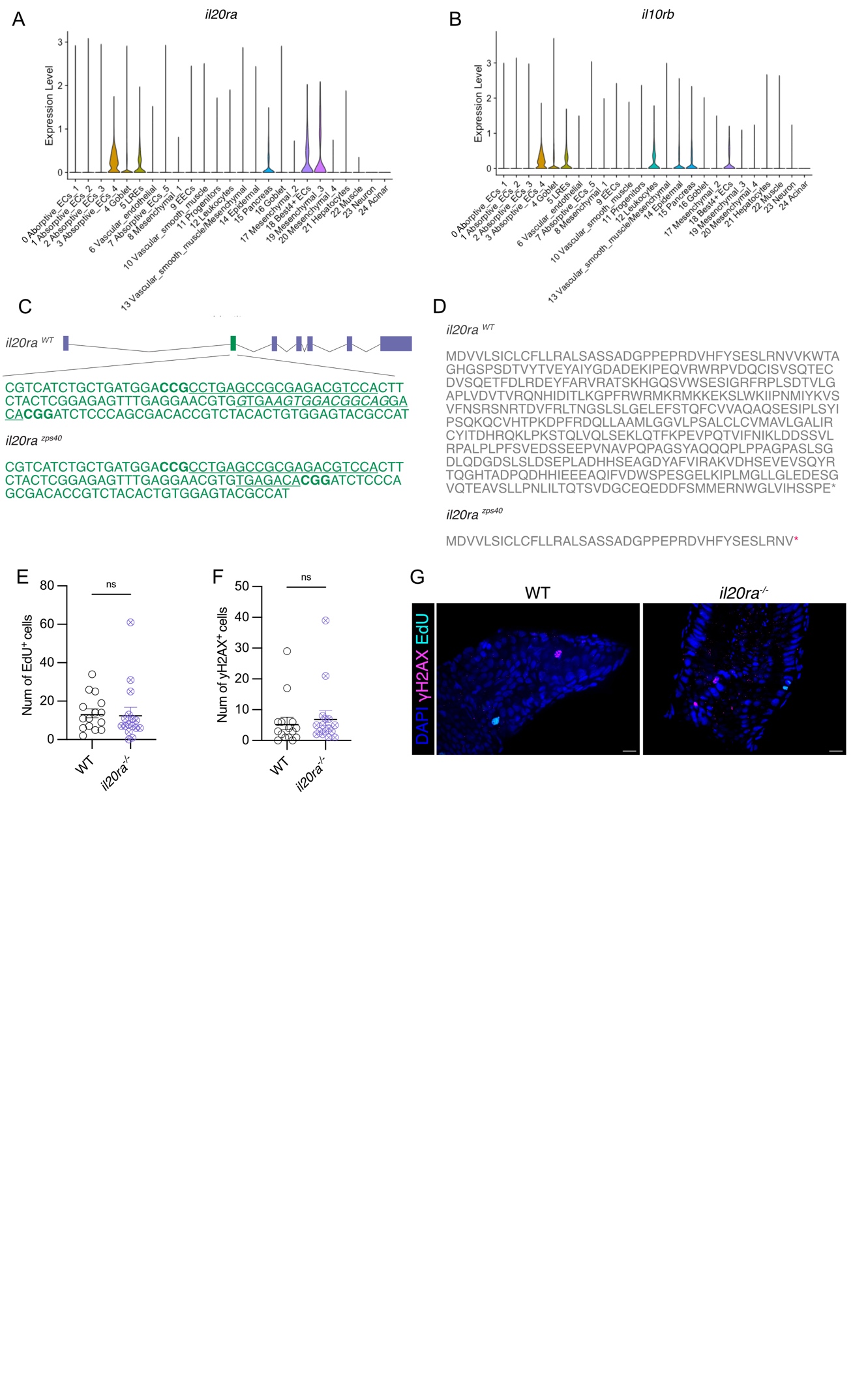

**Fig. S4: IL-26 regulates gut epithelial homeostasis in a receptor-independent manner.** Violin plot of *il20ra* (**A**) and *il10rb* (**B**) expression in the integrated scRNA-seq dataset of WT and *il26^-/-^* guts. (**C**) Schematics and DNA sequence of the WT and mutant zebrafish *il20ra* alleles. The single guide RNA (sgRNA) target sites are underlined. The protospacer adjacent motif (PAM) sequences are indicated in bold. The deleted nucleotides in *il20ra^zps40^* are highlighted in *italics*. This led to a deletion in the 2nd exon of the zebrafish *il20ra* gene (*il20ra^aps40^*). (**D**) The predicted protein sequence for *il20ra^zps40^* consisting of 40 amino acids. Quantification of EdU staining (**E**) and γH2AX staining (**F**) in WT and *il20ra^-/-^* 5 dpf larval guts. (**G**) Representative images of EdU and γH2AX staining in WT and *il20ra^-/-^* 5 dpf larval guts. Scale bars, 10 μm. Error bars show means ± SEM. Statistical significance was determined by Wilcoxon test (E and F).

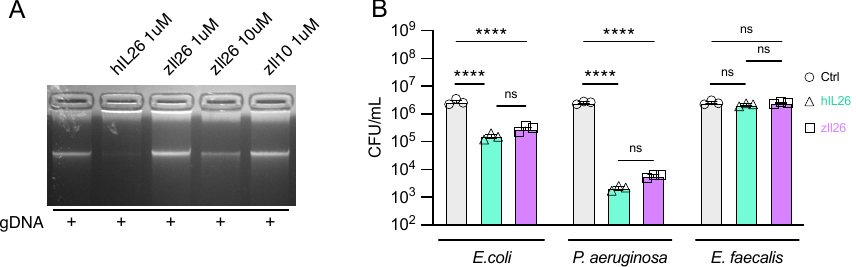

**Fig. S5: Analysis of IL-26 receptor-independent functions reveals conservation of IL-26 bactericidal activity in zebrafish.** (**A**) Gel migration assay of genomic DNA incubated with several cytokines. (**B**) Quantification of colony-forming units of different bacterial species after incubation with human or zebrafish IL-26 proteins at a concentration of 8 µM. Error bars show means ± SEM. Statistical significance was determined by 2way ANOVA (B).

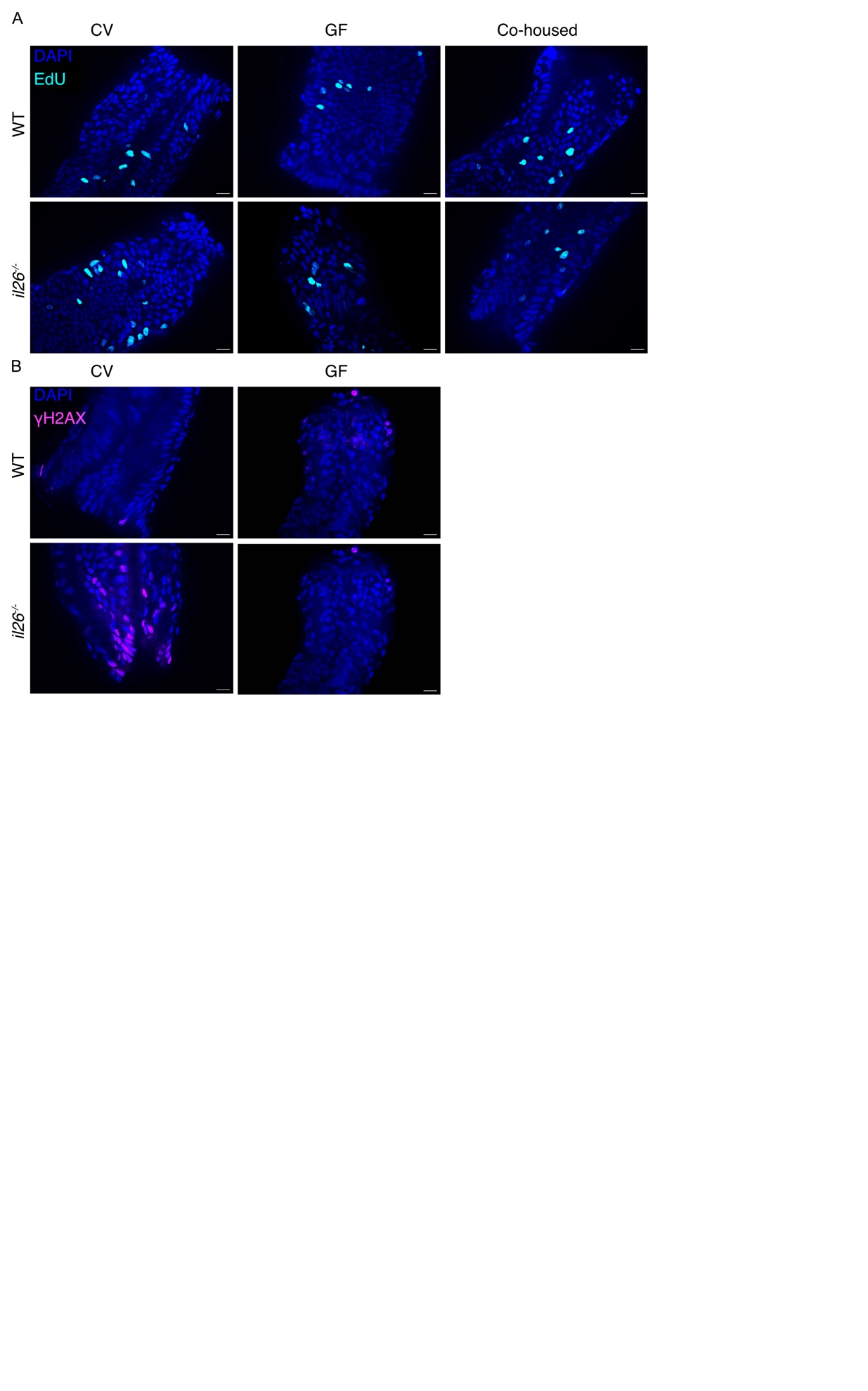

**Fig. S6: Immunostaining in WT and *il26^-/-^* larval guts reared CV, GF, or cohoused.** Representative images of EdU (**A**) and γH2AX staining (**B**) in WT and *il26^-/-^* larval guts reared CV, GF, or cohoused. Scale bars, 10 μm.

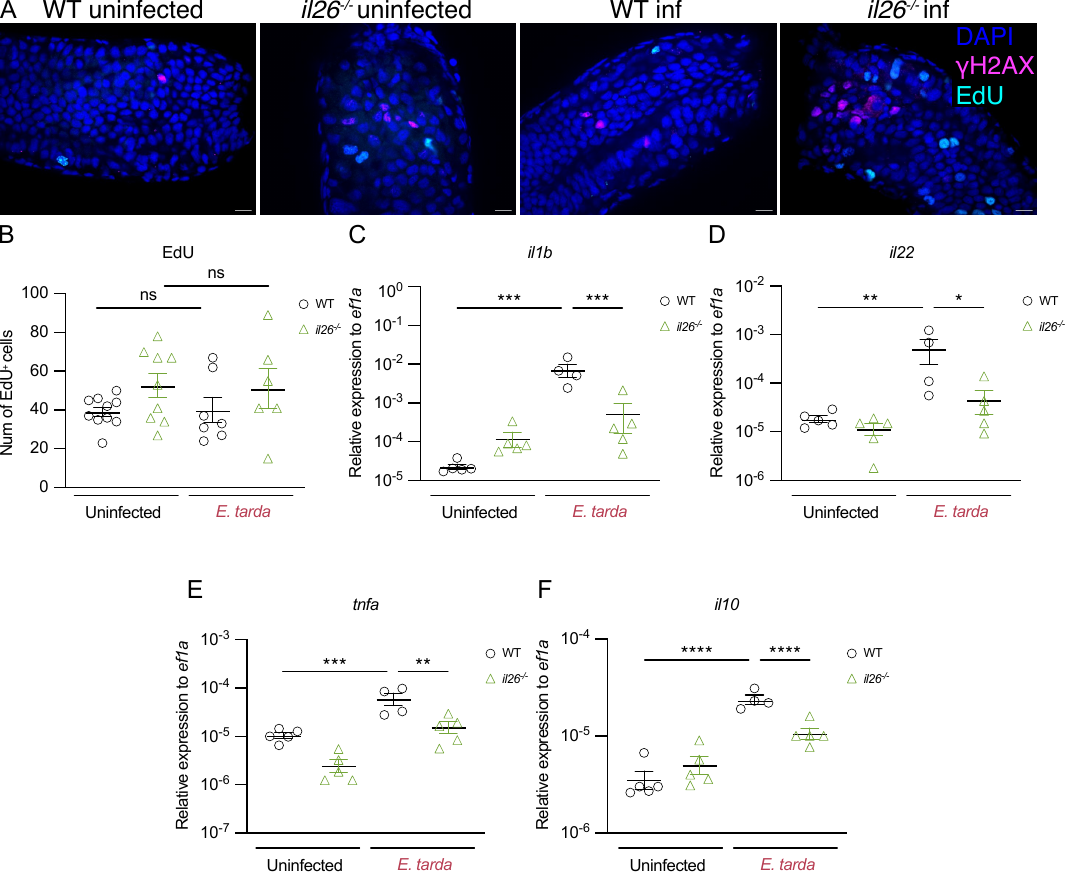

**Fig. S7: *il26^-/-^* larval guts mount an insufficient immune response upon *E. tarda* infection.** (**A**) Representative images of EdU and γH2AX staining in WT and *il26^-/-^* at 3 dpi. Scale bars, 10 μm. (**B**) Quantification of EdU staining in WT and *il26^-/-^* at 3 dpi. ) qRT-PCR analysis of *il1b* (**C**), *il22* (**D**), *tnfa* (**E**), *il10* (**F**) in dissected guts of WT and *il26^-/-^* at 3 dpi. Error bars show means ± SEM. Statistical significance was determined by one-way ANOVA (B-F). ns *P* > 0.05, **P* < 0.05, ***P* < 0.01, ****P* < 0.001, and *****P* < 0.0001.

Data S1. (separate file)

Survival rates of *il26^-/-^* fish at 3 months post-fertilization.

Data S2. (separate file)

Differentially expressed genes in 5 dpf *il26^-/-^* guts compared to WT controls.

Data S3. (separate file)

Differentially expressed genes in the guts of 5 dpf WT larvae injected with recombinant zebrafish IL-26 protein compared to BSA-injected controls.

Data S4. (separate file)

Marker genes identified from graph-based clustering of WT and *il26^-/-^* single-cell transcriptomes.

Data S5. (separate file)

Differentially expressed genes in the guts of 5 dpf GF *il26^-/-^* compared to GF WT controls.

Data S6. (separate file)

Taxon-based analysis of the 16S rRNA-sequencing from 5 dpf WT and *il26^-/-^* larvae.
